# SLiM 3: Forward genetic simulations beyond the Wright–Fisher model

**DOI:** 10.1101/418657

**Authors:** Benjamin C. Haller, Philipp W. Messer

## Abstract

With the desire to model population genetic processes under increasingly realistic scenarios, forward genetic simulations have become a critical part of the toolbox of modern evolutionary biology. The SLiM forward genetic simulation framework is one of the most powerful and widely used tools in this area. However, its foundation in the Wright–Fisher model has been found to pose an obstacle to implementing many types of models; it is difficult to adapt the Wright–Fisher model, with its many assumptions, to modeling ecologically realistic scenarios such as explicit space, overlapping generations, individual variation in reproduction, density-dependent population regulation, individual variation in dispersal or migration, local extinction and recolonization, mating between subpopulations, age structure, fitness-based survival and hard selection, emergent sex ratios, and so forth. In response to this need, we here introduce SLiM 3, which contains two key advancements aimed at abolishing these limitations. First, the new non-Wright–Fisher or “nonWF” model type provides a much more flexible foundation that allows the easy implementation of all of the above scenarios and many more. Second, SLiM 3 adds support for continuous space, including spatial interactions and spatial maps of environmental variables. We provide a conceptual overview of these new features, and present several example models to illustrate their use. These two key features allow SLiM 3 models to go beyond the Wright–Fisher model, opening up new horizons for forward genetic modeling.

## Introduction

Forward genetic simulations are playing an increasingly important role in evolutionary biology due to their ability to model a wide range of population genetic mechanisms and include a high level of ecological detail in the simulated scenario (Carvajal-Rodriguez, 2010; Yuan et al., 2012; Bank et al., 2014; Hoban, 2014; Thornton, 2014; Haller and Messer, 2017; Haller et al., 2018). The SLiM forward genetic simulation framework (Messer, 2013; Haller and Messer, 2017) has proved to be a powerful tool for this purpose, and arguably constitutes the most widely-used computational framework for implementing such simulations at the present time.

This popularity is based upon three key attributes of SLiM. First, it is highly scriptable, allowing the mechanics of the SLiM framework to be fundamentally modified and extended in many ways. At the same time, even fairly sophisticated models can often be expressed in a page of code or less, since all of the core simulation code is provided by SLiM, yielding tremendous benefits compared to writing simulations from scratch in a language such as C++. Second, SLiM includes a full-featured graphical modeling environment, SLiMgui, that makes interactive model development, visual debugging, and hands-on exploration easy, with large benefits throughout the modeling process (Grimm, 2002). And third, a great deal of work has been devoted to optimizing SLiM, making it run as efficiently as possible across a wide variety of simulation scenarios; these speed benefits are inherited for free by any model running within SLiM.

In our contact with users of SLiM, however, one category of questions has predominated: how can SLiM simulations be constructed that go beyond the standard Wright–Fisher or “WF” model (Fisher, 1922; Wright, 1931)? This model, which has provided the conceptual foundation for all previous versions of SLiM (Messer, 2013; Haller and Messer, 2017), is defined by a number of simplifying assumptions. For example, the model assumes that generations are non-overlapping and discrete, without any age structure or age-based differentiation among individuals. Another critical assumption of the model lies in the rules governing the generation of offspring from the parental population; in the standard WF model, the parents for each child in the next generation are drawn randomly from the previous generation with a probability proportional to each individual’s fitness. This makes it difficult to model variation in litter size, monogamous mating, and other such phenomena. Furthermore, since population size is an externally determined parameter in the WF model, it is often not clear how scenarios in which population size is an emergent variable – depending, for instance, on factors such as mean fitness, available habitat, and colonization history – should be accurately modeled in a WF framework.

Given the scope of the simplifying assumptions underlying the WF model, the desire among SLiM’s users to go beyond this model takes many forms, but they might be said to unify around the idea of more realistic spatial and ecological dynamics. For example, users have inquired whether it is possible to model the explicit movement of individuals over a continuous landscape, life cycles with overlapping generations, individual variation in reproduction, density-dependent population regulation, individual variation in dispersal or migration, local extinction and recolonization, mating between subpopulations, age structure, fitness-based survival and hard selection, emergent sex ratios, and more. Because SLiM 2 was already highly scriptable, and thus many of its internal dynamics could be modified through scripting, it was sometimes possible to work around the limitations inherited from the WF model; but those workarounds are often clumsy and laborious, and some types of models have simply proved difficult or impossible to implement in SLiM 2. Fundamentally, the Wright–Fisher model is not an ecological model, and so if we are to progress toward uniting genetics and evolutionary biology with ecology the need for a more flexible foundation is clear.

In response to this need, we here introduce SLiM 3, which contains two major advances squarely aimed at these limitations. First, in addition to the traditional Wright–Fisher or WF model type of previous SLiM versions, SLiM 3 supports a new non-Wright–Fisher or “nonWF” model type that provides much greater flexibility in how key processes such as mate choice and reproduction, migration, fitness evaluation, survival, population regulation, and other related areas are implemented, allowing the explicit linking of evolutionary dynamics with ecological patterns and processes. Second, in addition to support for discrete subpopulations connected by migration, SLiM 3 now supports models that occupy continuous spatial landscapes, including built-in support for spatial maps that describe environmental characteristics, and for local spatial interactions such as spatial competition and mate choice. (Support for spatial models was introduced in SLiM 2.3, in fact, but is previously unpublished).

SLiM 3 contains many other important additions as well. Most prominently, it adds support for “tree-sequence recording” (also called “pedigree recording”), a method of recording ancestry information in forward simulations (Kelleher et al., 2016; Kelleher et al., 2018). Tree-sequence recording can decrease simulation runtimes by orders of magnitude, by allowing neutral mutations to be overlaid efficiently after forward simulation has completed and by allowing neutral burn-in to be done extremely efficiently with “recapitation”, and it provides several other major benefits as well (Kelleher et al., 2018; Haller et al., 2018). SLiM 3’s support for tree-sequence recording is discussed further in Haller et al. (2018). Other important changes in SLiM 3 since SLiM 2.0 (the last published version) include many additions and improvements to the Eidos scripting language (Haller, 2016), many new methods provided by SLiM’s Eidos classes (Haller and Messer, 2016), many improvements to the SLiMgui graphical modeling environment, and a great deal of optimization work to make SLiM faster. A few more specific improvements are also worth mentioning: a new Individual class representing simulated individuals, support for a variable mutation rate along the chromosome, a new `recombination()` callback mechanism for modifying recombination breakpoints at an individual level, and VCF format output, among others (a complete change list may be found in the SLiM manual).

Here, however, we will focus on what we believe to be the most important new features in SLiM 3: nonWF models and continuous space, the features that enable users to go beyond the Wright–Fisher model. We will provide a conceptual overview of these features, and will demonstrate them with several examples.

### The nonWF Simulation Model

Perhaps the easiest way to understand nonWF models is by looking at how they differ from the standard WF model type. The most important differences are in the following broad areas:

- **Age structure.** In WF models, generations are discrete and non-overlapping; all individuals live for a single generation, during which they reproduce and then die. In nonWF models, by contrast, generations can be overlapping; individuals can live for multiple generations, until they die due to some cause (typically selection, old age, or bad luck). More fundamentally, the concept of a “generation” has been broadened. In nonWF models, each generation represents an opportunity to reproduce and/or die – a discretization of those events in time, providing a temporal structure to the model that could be based upon hours, days, seasons, or decades, but that is not necessarily related to the expected lifespan of individuals. Individuals in nonWF models have an age (measured in generations), the population thus has an age structure, and the model can implement whatever age-related behaviors are desired. The generation cycle in nonWF models is contrasted against that of WF models in Figure 1.

**Figure 1.**
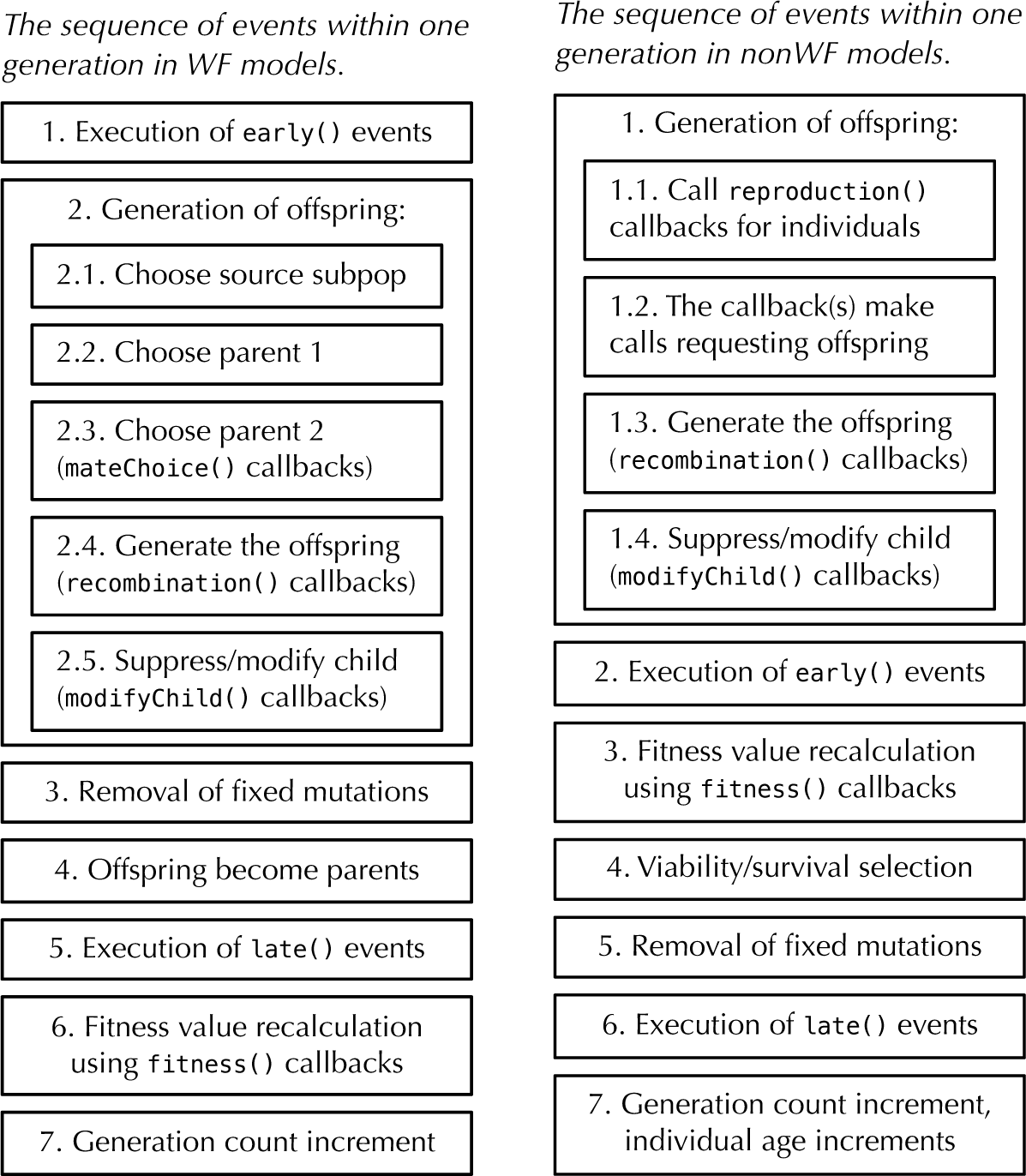
A comparison of the generation cycles in WF models (left) versus nonWF models (right). Note that nonWF models have a viability/survival selection phase, immediately after fitness value recalculation, whereas in WF models fitness influences mating success and there is no concept of mortality-based selection. Also, in WF models early() events come before offspring generation and late() events come after; in nonWF models, early() events come after offspring generation and late() events, by virtue of being at the end of the generation cycle, in effect come before offspring generation (when it occurs at the beginning of the next generation).
- **Offspring generation.** In WF models, offspring are generated by drawing parents from the individuals in the previous generation. The population size is a parameter of the model, determining how many offspring are to be generated in each generation; the selfing rate, cloning rate, and sex ratio are, similarly, population-level parameters. In nonWF models, by contrast, the script is much more directly in charge of the process of offspring generation; the script requests the generation of each offspring based upon individual state rather than population-level parameters. Calls are made from SLiM to `reproduction()` callbacks defined in the model script, and those callbacks determine which individuals reproduce, how they choose a mate or mates (if any), how many offspring they have, and so forth. The population size, selfing rate, cloning rate, and sex ratio are therefore no longer population-level parameters; instead, they are emergent properties of the model, consequences of the rules specified in script for the individual-based reproduction dynamics. The population size, for example, becomes the result of the balance between birth rates and death rates, often (but not necessarily) regulated by density-dependent viability selection implemented in the model.
- **Migration.** In WF models, migration between populations is modeled by specifying the fraction of offspring in a given target population that stem from parents in a given source population. Since this model of migration leads to offspring that occupy a different population from their parents, it most closely resembles a model of juvenile migration. In nonWF models, by contrast, migration is again handled more directly by the model script, which may call `takeMigrants()` to move individuals to a new population at any point in the generation cycle. This allows individuals to migrate as juveniles, as adults, or at multiple times during their life. This focus on individual-level migration, rather than population-level migration rates, allows for the probability that a given individual will migrate to depend much more flexibly upon individual-level state, enabling the modeling of a wide range of scenarios such as sex-dependent migration, habitat choice, or condition-dependent migration. In such models, the overall migration rate between two populations is again an emergent property that depends on the specific composition of the population and the migration rules specified in script, rather than on a population-level rate.
- **Fitness.** In WF models, fitness influences the probability that an individual will be chosen as a parent for a child in the next generation; there is no built-in concept of selection-induced mortality. Fitness is therefore relative, resulting in a model of so-called “soft” selection in which greater success for some individuals necessarily comes at the price of diminished success for others. The overall population size is not affected by selection, since it is a model parameter rather than an emergent property of the underlying evolutionary and ecological dynamics. In nonWF models, by contrast, fitness directly influences the probability of survival for each individual during each generation; individuals with low fitness are less likely to survive. Fitness is therefore absolute, and selection is “hard” in such a model by default; as a result, population size will vary naturally with mean population fitness (although this may be compensated for by density-dependent selection or fecundity). Of course one may still model the effects of genetics upon reproductive success or fecundity, in a `reproduction()` callback, if desired.

These differences can be summarized by saying that nonWF models are more individual-based, more script-controlled, more emergent, and therefore more biologically realistic. However, they are also often more complex in certain ways, primarily because of the need to implement a `reproduction()` callback and to introduce some explicit mechanism of population regulation. In effect, with the power to more precisely control reproduction and population regulation comes the responsibility to more explicitly think about and specify those phenomena. Populations can be regulated by any of a wide variety of mechanisms, from density-dependent fecundity to resource competition to predation to territorial behavior to natural disasters (Hixon et al., 2002; Begon et al., 2006). Any of these mechanisms can be implemented in a nonWF model, but it is not done for you as it is in a WF model; the user must decide what mechanism(s) of population regulation are desired and implement them in the model’s script.

To make this more clear, we will now look at some examples.

### Example 1: A simple nonWF model

Our first example is of a simple nonWF model that illustrates the basic structure and generation cycle involved. A `reproduction()` callback chooses a random mate and generates a single offspring. Density-dependent selection imposes a fitness cost upon individuals that depends on the population size, leading to an emergent population size that fluctuates stochastically around the carrying capacity of the model. Age structure is emergent too, with some individuals surviving through multiple generations (which might be thought of as years, if we assume one litter per year), while others perish as juveniles.

~~~
initialize() { initializeSLiMModelType("nonWF");
  defineConstant("K", 500); // carrying capacity
  initializeMutationType("m1", 0.5, "f", 0.0);
  m1.convertToSubstitution = T;
  initializeGenomicElementType("g1", m1, 1.0);
  initializeGenomicElement(g1, 0, 99999);
  initializeMutationRate(1e-7); initializeRecombinationRate(1e-8);
}
reproduction() {
  subpop.addCrossed(individual, p1.sampleIndividuals(1));
}
1 early()
{
  sim.addSubpop("p1", 10);
}
early() {
  p1.fitnessScaling = K / p1.individualCount;
}
2000 late() { sim.outputFixedMutations(); }
~~~

First of all, the script sets up the overall structure of the model in its initialize() callback. The model is declared to be a nonWF model with the call to `initializeSLiMModelType()`; this tells SLiM to use the nonWF generation cycle and so forth. Next, a constant `K` is defined, representing the nominal carrying capacity of the model as we will discuss further below. The remaining lines of the `initialize()` callback will be familiar to users of SLiM 2 as standard simulation configuration steps. They declare that the simulation uses only a single mutation type, representing neutral mutations; that it uses only a single genomic element type, representing genomic regions that undergo mutations of that mutation type; that the chromosome is of length 105 and is composed of a single such genomic element type; that mutations occur at a rate of 10^−7^ per base position per generation; and that recombination occurs at a rate of 10^−8^ per base position per generation. These concepts are discussed in more detail in the SLiM manual (Haller and Messer, 2016).

Second, the constant `K` is used in the `early()` event to calculate a density-dependent fitness effect that is set into `p1.fitnessScaling`, a new property. In SLiM 3, both individuals and subpopulations have `fitnessScaling` properties, and the final fitness value for each individual is calculated (multiplicatively) from the fitnessScaling of the subpopulation and the individual, as well as the effects (as in SLiM 2) of all mutations and `fitness()` callbacks. This provides a convenient way of altering fitness values, on either an individual or subpopulation level, without having to implement a `fitness()` callback. Here, it scales the absolute fitness of every individual so that the population will expand up to the carrying capacity. Importantly, fitness in this model, by default, specifies viability – the probability that an individual survives the viability/survival stage of the generation cycle. Note that the carrying capacity will not be met exactly in every generation; instead, the population size will fluctuate naturally around `K`, depending upon stochastic variation in fitness-based mortality. The formula in the `early()` event will result in a dynamic equilibrium around a population size of `K`. This is because if `p1.individualCount` is less than `K` every individual will be assigned a `fitnessScaling` value greater than 1.0, whereas if `p1.individualCount` is greater than `K` then `fitnessScaling` values will be less than 1.0. However, this scaling only results in equilibrium at size `K` if other fitness effects are not present; one might think of it as a “baseline carrying capacity”, from which the model might depart if other factors influencing survival or reproduction are present.

Third, a `reproduction()` callback has been defined. This callback is called once for each individual alive at the point of offspring generation, and is expected to generate offspring (if any) for the focal individual. Here, a mate is chosen at random from `p1` with `sampleIndividuals()`, and then biparental mating between the focal individual and the chosen mate is conducted by `addCrossed()`, adding the resulting offspring to the subpopulation of the focal individual. This code is entirely generalizable; multiple offspring could be generated, multiple mates could be chosen, and offspring could be generated through cloning or selfing instead of crossing.

The last point to note is that `convertToSubstitution` is set to `T` for `m1`, allowing fixed neutral mutations to be converted into Substitution objects by SLiM. In a WF model this would be automatic; the default is `convertToSubstitution=T`. In nonWF models, the default is `convertToSubstitution=F`. This is because since fitness is absolute, not relative, it would not be generally safe to remove fixed mutations from the simulation (but since m1 is neutral and has no indirect effects in the model, it is safe in this case). Telling SLiM that it can remove fixed mutations allows the model to run much faster.

Individuals in this model live until they happen to die (rather than being predestined to die at the end of each generation, as in a WF model); the age structure is emergent, determined only by birth and death rates. In the Eidos console in SLiMgui, we can see what age structure happens to emerge by executing `p1.individuals.age`. This reveals that, at the end of one randomly chosen test run, just one individual was 9 generations old, a few were 8 or 7, and about half the population was 1. This makes sense: in this model one new offspring is generated for every living individual in each generation, and then an equal probability of death for every individual in each generation, combined with the density-dependent selection and the specified carrying capacity, should produce the observed age distribution.

### Example 2: Age structure and monogamy

This slightly more advanced nonWF model illustrates two features. The first is how the age structure of the model can be controlled, with a minimum age for reproduction and varying survival rates depending upon age. The second is how to implement monogamy with litter size drawn from a distribution.

~~~
initialize() {
initializeSLiMModelType("nonWF");
defineConstant("K", 500); // carrying capacity
initializeMutationType("m1", 0.5, "f", 0.0);
m1.convertToSubstitution = T;
initializeGenomicElementType("g1", m1, 1.0);
initializeGenomicElement(g1, 0, 99999);
initializeMutationRate(1e-7);
initializeRecombinationRate(1e-8);
}
reproduction() {
if (individual.age >= 3)
{
mate = p1.sampleIndividuals(1, minAge=3);
litterSize = rpois(1, 2);
for (i in seqLen(litterSize))
subpop.addCrossed(individual, mate);
}
}
1 early() {
sim.addSubpop("p1", 10);
}
early() {
p1.fitnessScaling = K / p1.individualCount;
inds = p1.individuals;
inds.fitnessScaling = ifelse(inds.age < 5, 1.0, 0.1);
}
2000 late() { sim.outputFixedMutations(); }
~~~

The `initialize()` callback is unchanged. The `reproduction()` callback specifies that only individuals of age `3` or older can reproduce, and the same requirement is enforced upon the chosen mate with `sampleIndividuals()`. A litter size is drawn from a Poisson distribution with mean `2`, and then all of the offspring in the litter are generated by calls to `addCrossed()`. Monogamy is modeled here, but any mating scheme could be enforced through the appropriate script.

The other difference is in the `early()` event, where an individual fitness effect, based upon age, is applied to the `fitnessScaling` property of the individuals. Individuals of age less than `5` receive no individual scaling (but still feel the fitness effects of density, mutations, etc.), whereas individuals of age `5` or greater receive a multiplicative fitness penalty of `0.1`; it is still possible to live to a ripe old age, but much less likely than in the previous model. Individuals in this model typically live to be `5` or fewer generations old, although outliers are possible. This age distribution was not specified as a model parameter, however; it is an emergent consequence of the mechanistic biological details specified, such as the minimum age of reproduction, the mean litter size, and the fitness effect of age.

Population size is also emergent in this model; we apply a “baseline carrying capacity” using a value of 500 for `K`, as in Example 1, but because fitness values are also influenced by age-based mortality, the actual equilibrium population size is a little lower than `K` (about 480). This is as it should be; the model is doing precisely what we have told it to do in script. If a different model of population regulation were desired, that could be implemented in the script instead; all of the power over population regulation is in the hands of the modeler with nonWF models. Of course, this tremendous flexibility comes at a price: the mechanism of population regulation must be carefully planned in order to prevent unrealistic population dynamics, such as runaway growth or unintended population extinction.

These same concepts apply similarly to selfing rate, cloning rate, sex ratio (in sexual models), and migration rates: in a nonWF model, the mechanistic biological details are specified, and the behavior of the model is emergent from that specification (various recipes illustrating these possibilities are shown in the SLiM manual). Let us now move on to the topic of continuous space, the other major way in which SLiM 3 facilitates going beyond the Wright–Fisher model.

### Continuous Space and Spatial Interactions

Continuous-space models in SLiM 3 are quite straightforward at the conceptual level.

Continuous space is enabled with a call to `initializeSLiMOptions()` that provides a dimensionality: `“x”` for one spatial dimension, `"xy"` for two, or `"xyz"` for three; we will focus here on 2D models since that is probably the most common case. Individuals then have properties representing their `x` and `y` coordinates in the continuous 2D space, which can be accessed and set. The spatial boundaries of each subpopulation can be configured by the user; by default, the landscape will span the interval [0,1] in each dimension. Setting individual positions is the responsibility of the model, and the model determines what use, if any, is made of those positions; there is no automatic consequence of spatiality upon model dynamics. However, since there are common ways in which models often want spatiality to influence dynamics, two additional facilities are provided: interaction types, and spatial maps.

Interaction types are supported with a new Eidos class, `InteractionType`. An interaction type is defined with a call to `initializeInteractionType()`, and specifies two things: a distance metric that determines the interaction distance between two individuals, and an interaction formula that determines how the strength of interaction between two individuals varies with the distance between them. Once an interaction type is set up and evaluated, spatial queries can be made: what are the *n* closest neighbors to a given individual, what is the strength of interaction between individuals *i* and *j*, what is the total interaction strength exerted upon individual *i* by all other individuals in its subpopulation, and so forth. These queries are handled internally by highly optimized data structures such as *k*-d trees (Bentley, 1975) and sparse arrays (Tewarson, 1973), but those details are entirely hidden by SLiM, providing a way of implementing spatial interactions such as spatial competition and spatial mate choice that is both simple and fast.

Spatial maps are not represented with a separate class; instead, they are attached to subpopulations. A new spatial map can be defined with a call to the `defineSpatialMap()` method, and the value of a particular spatial map at a given point can then be queried with `spatialMapValue()`. Any number of spatial maps may be attached to a subpopulation; multiple maps are distinguished from each other by name. Each map defines a grid of values (of any resolution) that is superimposed across the spatial bounds of the subpopulation, either with or without interpolation of values between grid points. The scale of the map values, and the meaning attached to them, is entirely up to the model to define. One map might define elevation across the landscape, another temperature, and each of those maps might have consequences for survival, or fecundity, or movement, or any other aspect of the model.

Since much of this may seem rather abstract, we will now look at two concrete examples.

#### Example 3: A spatial mate choice model

We will start with a spatial model of individuals living on a homogeneous two-dimensional landscape. The population size is regulated through local competition, using an `InteractionType object, i1`, to tally up its effects on each individual. Individuals prefer nearby individuals as mates, using another `InteractionType object, i2`, to find a mate. Each offspring is generated from an independently chosen mate, and lands near its first parent in space. This model uses “periodic” boundary conditions; the landscape wraps around both horizontally and vertically, providing a seamless toroidal space that avoids any boundary effects. A beneficial mutation is introduced in generation `1000`, which might sweep through the population or might be lost. The model:

~~~
initialize() {
initializeSLiMModelType("nonWF");
initializeSLiMOptions(dimensionality="xy", periodicity="xy"); defineConstant("K", 2000); // carrying-capacity density defineConstant("S", 0.05); // sigma_S, the spatial interaction width
initializeMutationType("m1", 0.5, "f", 0.0); // neutral m1.convertToSubstitution = T;
initializeMutationType("m2", 0.5, "f", 0.1); // beneficial
initializeGenomicElementType("g1", m1, 1.0);
initializeGenomicElement(g1, 0, 99999);
initializeMutationRate(1e-7); initializeRecombinationRate(1e-8);
initializeInteractionType(1, "xy", reciprocal=T, maxDistance=S * 3); i1.setInteractionFunction("n", 1.0, S);
initializeInteractionType(2, "xy", reciprocal=T, maxDistance=0.02); i2.setInteractionFunction("n", 1.0, 0.01);
}
reproduction() {
mate = i2.drawByStrength(individual, 1);
for (i in seqLen(rpois(1, 0.1)))
{
if (mate.size())
offspring = subpop.addCrossed(individual, mate);
else
offspring = subpop.addSelfed(individual);
pos = individual.spatialPosition + rnorm(2, 0, 0.02);
offspring.setSpatialPosition(p1.pointPeriodic(pos));
}
}
1 early() {
sim.addSubpop("p1", 1);
p1.individuals.setSpatialPosition(p1.pointUniform(p1.individualCount));
}
1000 early() {
sample(p1.genomes, 1).addNewDrawnMutation(m2, 20000);
}
early() {
i1.evaluate();
inds = p1.individuals;
competition = (i1.totalOfNeighborStrengths(inds) + 1) / (2 * PI * S^2); inds.fitnessScaling = K / competition;
mut = sim.mutationsOfType(m2); if (mut.size())
inds.color = ifelse(inds.containsMutations(mut), "blue", "");
}
late()
{
i2.evaluate();
}
10000 late() { sim.outputFixedMutations();
}
~~~

This model is complex enough that we can’t go into great detail on it, but we can sketch its broad design. In the `initialize()` callback we specify a nonWF model with dimensionality `"xy"`, with both dimensions being periodic. A mutation type is defined for neutral mutations, and another for the beneficial mutation that will be introduced. The model’s genetic structure is specified as usual. Finally, two interaction types are set up; i1 is for spatial competition, using a Gaussian kernel of width S (with a maximum distance of `S*3`), whereas `i2` is for spatial mate choice, using a Gaussian kernel of width `0.01` (maximum distance `0.02`).

In the `reproduction()` callback the focal individual draws a mate from within the maximum interaction distance, weighted by interaction strength so that nearer individuals are more likely to be chosen, using `i2`. A litter is then generated, of a size drawn from a Poisson distribution, using crossing if a mate was found or selfing if not. The position of each offspring lands near its first parent, with accounting for the periodic boundaries of the model.

The next event creates an initial population of a single individual, and sets its initial position randomly. Selfing is thus essential for this model to first get started; but a larger initial population size may be used instead, of course. In generation `1000` a new `m2` mutation is introduced into a randomly chosen genome by the next event. The following `early()` event runs in every generation, and calculates the effect of local density upon the fitness of each individual using `i1` (the interactions of which are evaluated just prior to its use); the mathematical details of this type of local density-dependent regulation are explained in the SLiM manual (Haller and Messer, 2016). This local density dependence generates realistic spatial population dynamics, such as preventing excessive clustering and ensuring that individual fitness is influenced only by nearby individuals. This event also colors carriers of the beneficial mutation blue, for illustration purposes. Finally, `i2` interactions are evaluated in a `late()` event, just in time for them to be used by `reproduction()` callbacks at the beginning of the next generation; and then some token output is generated in generation `10000`.

If the beneficial mutation is not lost early on, it will typically sweep across the population slowly, with a clear spatial pattern in its spread, as shown in Figure 2. The spread is rather slow in this model (compared to a panmictic WF population) partly because it is limited by the dispersal of individuals in this spatial model, and partly because individuals in this model tend to be quite long-lived (with a mean age of approximately `10`).

**Figure 2.**
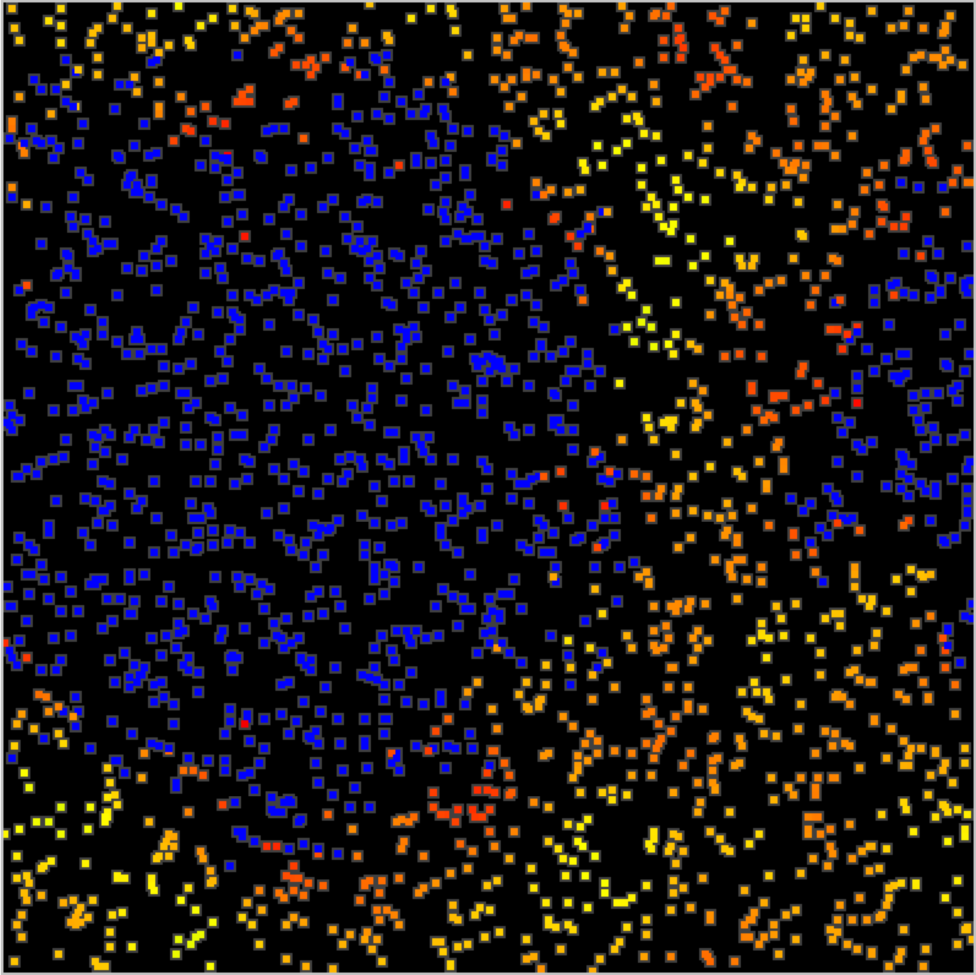
A snapshot of Example 3 at generation 1300, showing the spatial sweep of a mutation (blue) that was introduced in generation 1000. Spatial boundaries in this model are periodic, as seen in the way the sweep bridges the far left and far right. Individuals that do not possess the sweep mutation are colored to indicate their fitness, from yellow (neutral) to red (low); this variation in fitness is due to the effects of spatial competition, which regulates the population density. Only survivors after the selection stage of the generation cycle are visible, explaining what might otherwise appear to be incongruous fitness values.

#### Example 4: Spatial recolonization on a heterogeneous landscape

In this example, we introduce a spatial map that describes the habitability of the landscape. Areas of low habitability decrease the fitness of the individuals that occupy them, thereby decreasing the local population density. Occasional “disasters” will wipe out all individuals in a randomly chosen area of the map, which will lead to subsequent recolonization of the vacant areas in a wave limited by the landscape’s habitability. The model:

~~~
initialize() {
initializeSLiMModelType("nonWF");
initializeSLiMOptions(dimensionality="xy");
defineConstant("K", 2000); // carrying-capacity density defineConstant("S", 0.1); // sigma_S, the spatial interaction width
initializeMutationType("m1", 0.5, "f", 0.0); // neutral m1.convertToSubstitution = T; initializeGenomicElementType("g1", m1, 1.0);
initializeGenomicElement(g1, 0, 99999); initializeMutationRate(1e-7); initializeRecombinationRate(1e-8);
initializeInteractionType(1, "xy", reciprocal=T, maxDistance=S * 3); i1.setInteractionFunction("n", 1.0, S); initializeInteractionType(2, "xy", reciprocal=T, maxDistance=0.1);
}
reproduction() {
mate = i2.nearestNeighbors(individual, 1);
if (mate.size() & (runif(1) < 0.1)) {
offspring = subpop.addCrossed(individual, mate);
do pos = individual.spatialPosition + rnorm(2, 0, 0.02); while (!p1.pointInBounds(pos)); offspring.setSpatialPosition(pos);
}
}
1 early() { sim.addSubpop("p1", 100);
p1.individuals.setSpatialPosition(p1.pointUniform(p1.individualCount)); p1.defineSpatialMap("hab", "xy", c(10,10), runif(100, min=0, max=K),
interpolate=T, valueRange=c(0.0, K), colors=c("black", "blue"));
}
early() {
inds = p1.individuals; for (ind in inds)
ind.fitnessScaling = p1.spatialMapValue("hab", ind.spatialPosition);
i1.evaluate();
competition = (i1.totalOfNeighborStrengths(inds) + 1) / (2 * PI * S^2); inds.fitnessScaling = inds.fitnessScaling / competition;
if (runif(1) < 0.001) {
center = p1.pointUniform(1);
hit = inds[i1.distanceToPoint(inds, center) < 0.4]; hit.fitnessScaling = 0.0;
catn(sim.generation + ": BOOM!");
}
}
late()
{
i2.evaluate();
}
10000 late() { sim.outputFixedMutations();
}
~~~

The `initialize()` callback is similar to the previous model, but without periodic boundaries or beneficial mutations. The `reproduction()` callback now simply chooses the nearest individual as a mate, within the maximum distance of `i2`; `i2`’s kernel is unused and so was omitted. If a mate is found, there is a 10% chance of generating an offspring through crossing. “Reprising” boundaries are used here, meaning that new candidate positions for the offspring are drawn until a position within bounds is obtained.

The population initialization now sets up a spatial map, named `"hab"`, generated by 100 random values forming a 10×10 grid. Random values are between `θ` and `K`; the local habitability value from this map will take the place of the so-called “carrying-capacity density” from the previous model. The spatial map is interpolated, and we tell SLiM – purely for purposes of display in SLiMgui – that values from `0.0` to `K` should be given colors from black to blue. The next `early()` event then uses this spatial map to look up the local carrying-capacity density for each individual from the spatial map, and competition scales with that value rather than with a global carrying-capacity density. Finally, with a probability of 0.001 in each generation, disasters occur: a random point is chosen, and all individuals within a radius of 0.4 of the epicenter are given a `fitnessScaling` value of `0.0`, which is lethal.

Recolonization of disaster areas will occur by spatial spread from the surviving areas. However, dispersal will be largely restricted to high-habitability corridors, preventing recolonization from some areas that are nearby but disconnected. A snapshot of these dynamics soon after a disaster is shown in Figure 3.

**Figure 3.**
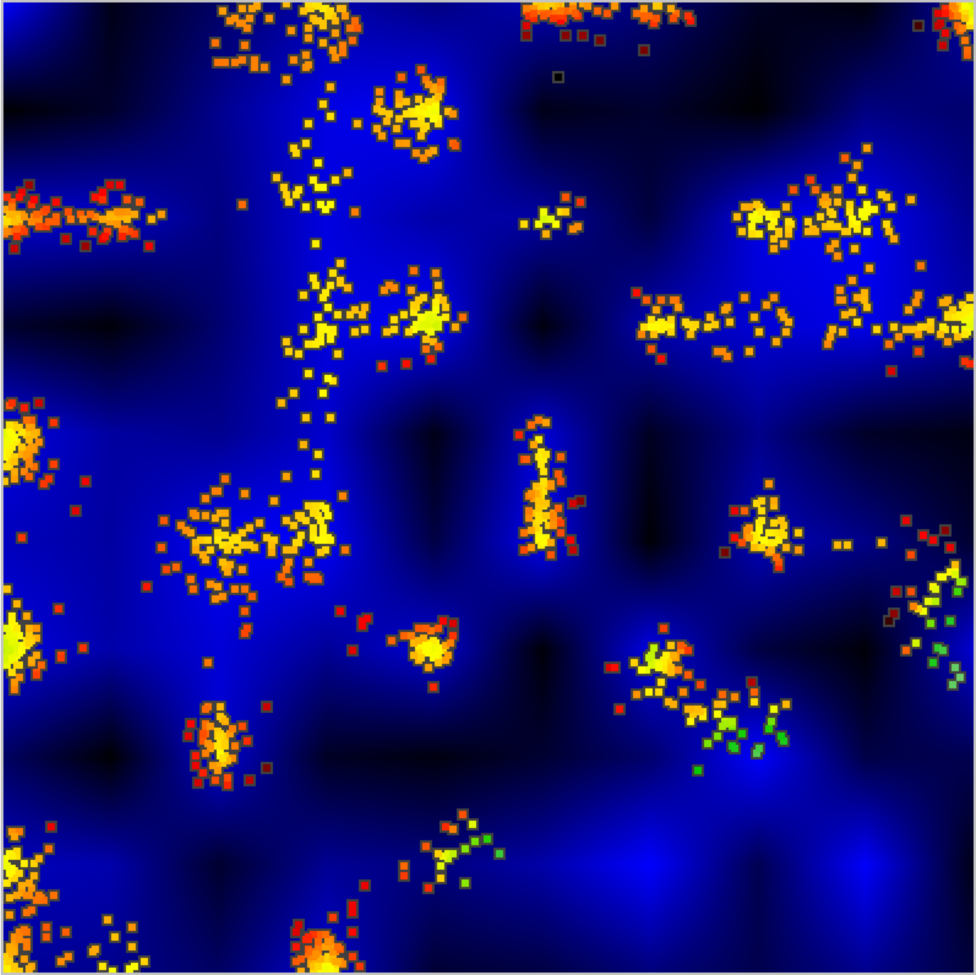
A snapshot of Example 4 at generation 657, after a disaster in generation 361 wiped out the individuals at lower right. Habitability, defined by a random spatial map with interpolation, is shown in background shades from black (uninhabitable) to blue (fully habitable). Recolonization of the disaster area is proceeding on three fronts, through corridors of relatively high habitability. Individuals at the three recolonization fronts are green, indicating their exceptionally high fitness because they are experiencing little competition from other nearby individuals.

Spatial maps can represent anything: real elevation values from an empirical study area, or levels of an environmental contaminant, or even calculated values related to the dynamics of the model, such as the mean age of individuals in the local area. Multiple spatial maps can be defined if desired; each is accessed by name. The possibilities are quite open-ended, and many more examples are shown in the SLiM manual (Haller and Messer, 2016).

## Discussion

We have presented SLiM 3, a new major release of the SLiM forward genetic simulation framework. SLiM 3 provides many improvements over previous versions of SLiM, which are described in detail in the comprehensive documentation. Here we have focused on the two major features that enable SLiM 3 models to go beyond the limitations and assumptions of the Wright– Fisher model upon which all previous versions were based: the nonWF model type, and support for continuous space.

The nonWF model type affords the model control over each individual mating event. This makes it easy to control model characteristics such as mate-choice behavior, fecundity, and individual variation in reproduction. In nonWF models, fitness influences survival, not mating probability, by default; this is typically more biologically realistic, and allows more natural population dynamics. Other important model features that are relevant for realistic ecology, such as overlapping generations, age structure, and more realistic migration/dispersal behavior, also emerge naturally in this design. However, the option to construct a WF model, as in previous SLiM versions, remains; this can be useful particularly when one wishes to compare a forward simulation model to an analytical model based upon Wright–Fisher assumptions.

Similarly, SLiM 3 provides the option of incorporating continuous space into a model, but models of discrete subpopulations connected by migration are also still supported. When continuous space is enabled, SLiM 3 provides a variety of useful tools for spatial modeling, such as spatial maps, which can define landscape characteristics that influence model dynamics, and a spatial interaction engine that can efficiently calculate interaction strengths between individuals and find nearby neighbors of an individual. SLiMgui also provides helpful visualizations for such models, making it easy to observe the dynamics that emerge from spatiality.

It should be emphasized that these features really dovetail with each other; in particular, ecologically realistic models involving continuous space should almost always be nonWF models. This is because the WF model imposes global population regulation upon the simulation; an overall size is set for each subpopulation, such that if density increases in one area of space (due to an immigration event, for example), absolute fitness will effectively decrease across the whole landscape. It is possible to compensate for this with appropriate fitness scaling, but it becomes quite complex if there is variation in local carrying capacity, immigration and emigration events, variation in fecundity, etc.; the externally imposed population size of WF models is simply not designed to accommodate locally-determined population density. The emergent population size and density in nonWF models, on the other hand, automatically accounts for whatever factors influence birth and death in the model. This is the reason that both of the continuous-space examples presented here are nonWF models; the influence upon local density of the sweep of a beneficial mutation in Example 3, or of the occasional disasters of Example 4, would be extremely difficult to model in a WF framework.

Many other new features of SLiM 3 have not been substantially discussed here. We urge all users to read about tree-sequence recording, which we believe to be a revolutionary new method that will considerably extend the horizon of what is possible in forward simulation (Haller et al., 2018). The SLiM manual (Haller and Messer, 2016) now contains recipes and reference documentation for other new features, and the Eidos manual (Haller, 2016) now documents new additions to the Eidos language. It is worth noting particularly that a great deal of optimization work has gone into SLiM 3, and it is generally much faster than previous versions – especially for large models with long chromosomes, which can be orders of magnitude faster than in previous versions.

SLiM 3 is free, licensed under the GNU GPL, and is available on GitHub. Most users, however, will wish to download the release version from https://messerlab.org/slim/; the extensive manuals, with many examples, can be downloaded from the same website. We also encourage SLiM users to subscribe to the slim-discuss list at http://bit.ly/slim-discuss, where new versions are announced and users can ask questions and get help. The features that we have focused on here, nonWF models and continuous space, will enable many modeling scenarios that would have been difficult or impossible to model previously. We hope this will open up new frontiers in evolutionary modeling, allowing both applied and theoretical research to go beyond the Wright–Fisher model.

## Acknowledgements

We thank Jared Galloway, Jerome Kelleher, and Peter Ralph for their work on the tree-sequence recording feature of SLiM 3. Thanks also to Jorge Amaya, Bill Amos, Jaime Ashander, Hannes Becher, Emma Berdan, Yoann Buoro, Deborah Charlesworth, Jean Cury, A.P. Jason de Koning, Emily Dennis, Jordan Rohmeyer Dherby, Jared Galloway, Jesse Garcia, Kimberley Gilbert, Alexandre Harris, Rebecca Harris, Ding He, Kathryn Hodgins, Christian Huber, Melissa Jane Hubisz, Jacob Malte Jensen, Jerome Kelleher, Bhavin Khatri, Bernard Kim, Athanasios Kousathanas, Benjamin Laenen, Stefan Laurent, Eugenio Lopez, Kathleen Lotterhos, Mikhail Matz, Rupert Mazzucco, Maéva Mollion, Miguel Navascués, Greg Owens, Denis Pierron, Peter Ralph, David Rinker, Murillo Fernando Rodrigues, Andrew Sackman, Aaron Sams, Kieran Samuk, Onuralp Söylemez, Kevin Thornton, Robert Unckless, Christos Vlachos, Silu Wang, Aaron Wolf, Justin Yeh, members of the Messer lab, and the stackoverflow community for their valuable comments. Thanks to Peter Ralph for helpful comments on a previous version of this manuscript. This work was supported by funds from the College of Agriculture and Life Sciences at Cornell University, New Zealand’s Predator Free 2050 program (grant number SS/ 05/01), and the National Institutes of Health (grant number R01GM127418) to P.W.M.

## References

Bank, C., Ewing, G.B., Ferrer-Admettla, A., Foll, M., Jensen, J.D. (2014). Thinking too positive? Revisiting current methods of population genetic selection inference. Trends in Genetics 30(12), 540–546.

Begon, M., Townsend, C.R., and Harper, J.L. (2006). Ecology: From Individuals to Ecosystems. Hoboken, NJ: Wiley-Blackwell. 750 pp.

Bentley, J.L. (1975). Multidimensional binary search trees used for associative searching. Communications of the ACM 18(9), 509–517.

Carvajal-Rodriguez, A. (2010). Simulation of genes and genomes forward in time. Current Genomics 11(1), 58–61.

Fisher, R.A. (1922). On the dominance ratio. Proceedings of the Royal Society of Edinburgh 42, 321–341.

Grimm, V. (2002). Visual debugging: A way of analyzing, understanding and communicating bottom-up simulation models in ecology. Natural Resource Modeling 15(1), 23–38.

Haller, B.C. (2016). Eidos: A Simple Scripting Language. URL: http://benhaller.com/slim/Eidos_Manual.pdf

Haller, B.C., and Messer, P. W. (2016). SLiM: An Evolutionary Simulation Framework. URL: http://benhaller.com/slim/SLiM_Manual.pdf

Haller, B.C., and Messer, P.W. (2017). SLiM 2: Flexible, interactive forward genetic simulations. Molecular Biology and Evolution 34(1), 230–240. DOI:http://dx.doi.org/10.1093/molbev/msw211

Haller, B.C., Galloway, J., Kelleher, J., Messer, P.W., and Ralph, P.L. (2018). Tree-sequence recording in SLiM opens new horizons for forward-time simulation of whole genomes. bioRxiv, DOI:https://doi.org/10.1101/407783

Hixon, M.A., Pacala, S.W., and Sandin, S.A. (2002). Population regulation: Historical context and contemporary challenges of open vs. closed systems. Ecology 83(6), 1490–1508.

Hoban, S. (2014). An overview of the utility of population simulation software in molecular ecology. Molecular Ecology 23(10), 2383–2401.

Kelleher, J, Etheridge, A.M., and McVean, G. (2016). Efficient coalescent simulation and genealogical analysis for large sample sizes. PLoS Computational Biology 12(5): e1004842.

Kelleher, J., Thornton, K.R., Ashander, J., and Ralph, P.L. (2018). Efficient pedigree recording for fast population genetics simulation. bioRxiv, DOI:http://dx.doi.org/10.1101/248500

Messer, P.W. (2013). SLiM: Simulating evolution with selection and linkage. Genetics 194(4), 1037–1039.

Tewarson, R.P. (1973). Sparse Matrices. New York, NY: Academic Press. 159 pp.

Thornton, K.R. (2014). A C++ template library for efficient forward-time population genetic simulation of large populations. Genetics 198(1), 157–166.

Wright, S. (1931). Evolution in Mendelian populations. Genetics 16, 97–159.

Yuan, X., Miller, D.J., Zhang, J., Herrington, D., and Wang, Y. (2012). An overview of population genetic data simulation. Journal of Computational Biology 19(1), 42–54.

